# Transcranial Magnetic Stimulation Differentiates Egocentric vs. Allocentric Network Hubs for Memory-Guided Reach

**DOI:** 10.64898/2026.07.29.741451

**Authors:** Lina Musa, Gaelle Nsamba Luabeya, Brando Sheldrick, Ali Rezaei, Saihong Sun, Xiaogang Yan, Michael Vesia, J. D. Crawford

## Abstract

Previous patient and imaging studies suggest that dorsal and ventral visual stream mechanisms contribute selectively to egocentric (eye-centred) and allocentric (landmark-centred) goal-directed behaviours, respectively. However, causal evidence for this ego-allocentric dissociation in healthy brains is lacking. Based on a recent neuroimaging study, we hypothesized that perturbations to dorsal stream network hubs would disrupt both egocentric and allocentric reaches, whereas perturbations to ventral hubs would affect only allocentric reaches. Sixteen participants viewed a briefly presented target in the presence of a landmark that later shifted after a memory interval. Participants were instructed to 1) ignore the landmark and reach toward / opposite the original target location (egocentric task) or 2) reach toward the target’s location fixed relative to the shifted landmark (allocentric task). Transcranial magnetic stimulation (TMS) was applied during the memory period, spanning the landmark shift, at two occipital sites, based on the network hub coordinates from our previous study and individual brain anatomy. As predicted, superior occipital TMS impaired reaction time, accuracy, and precision in both tasks, whereas inferior occipital TMS only impaired allocentric performance. Further, spatial asymmetries in the resulting error patterns were influenced by TMS site and task: superior occipital TMS influenced asymmetries more strongly in the egocentric task, whereas inferior occipital TMS only influenced spatial asymmetries in the allocentric task. Finally, a functional network analysis revealed a widespread influence of TMS on network-behavior relations and another dissociation: greater influence of superior occipital TMS on the egocentric task and of inferior occipital TMS on the allocentric task. These results establish specific causal relationships between occipital hubs, widespread cortical networks, and reach behavior, confirming a default dorsal egocentric transformation for visually aimed reaches, and a role for the ventral stream for landmark-centred actions.

## Introduction

Successful interactions with our surroundings depend on the ability to encode, retain, and transform spatial information into goal-directed actions. Visual target locations can be represented either in egocentric (relative to the observer) or allocentric (relative to visual landmarks) frames of reference [1–3]. These mechanisms can be dissociated by instructing participants to rely on one cue or the other (e.g., [4,5]). Using such tasks, human neuroimaging / neuropsychology experiments suggest that activity / damage within different cortical ‘streams’ correlates differentially with egocentric vs. landmark-centred aiming movements [6–8]. However, direct causal evidence for this dissociation in healthy individuals has not been established.

The idea that the visual system is divided into two streams was originally based on the spatial tuning of neurons recorded in animals [9] but was later refined by neuropsychological experiments in humans [10,11]. Specifically, damage to the ‘dorsal stream’ (from occipital cortex through posterior parietal cortex) is associated with deficits in egocentric, action-oriented transformations, whereas damage to the ventral stream (from occipital to temporal cortex) is associated with deficits in object- and landmark-centered representations for perception [10,11,7]. More recent neuroimaging studies confirmed a dorsal-ventral / ego-allocentric dissociation and localized these processes to specific anatomic sites. Specifically, dorsal occipital / parietal cortex is activated and spatially tuned for egocentric reaches, whereas inferior occipital / temporal cortex is activated and tuned for landmark centred reaches [12,4].

Under natural conditions, ego- and allocentric cues are combined for optimal behavior [13], with the weighting dependent on various factors, including visual conditions [14,15] and memory delay [16,17]. This suggests that some mechanism flexibly integrates ventral / allocentric signals into the dorsal parietal-frontal stream, where egocentric transformations from gaze-centered visual signals to body-centred motor commands predominate [18,19]. Neuroimaging and physiological experiments suggest this could occur as early as parietal or as late as prefrontal cortex [20,21].

Recently, Musa et al. [8] analyzed functional connectivity (temporal correlations in brain activity) using the same dataset recorded by Chen et al. (2014), employing graph theory [22]. We identified two ‘hubs’ (nodes with highly distributed, strong inter-node correlations) within the visual peripheral network [8]: a superior occipital hub for both the egocentric and allocentric reaching, and an inferior occipital hub for allocentric reaching. Further, modularity analysis revealed that dorsal and ventral stream modules (clusters of correlated nodes) were separated during egocentric reaches, but joined during landmark-centred reaches, suggesting task-dependent ventral-dorsal integration [8].

Overall, these findings suggest that the dorsal visual stream provides a default egocentric transformation for visually guided action, and the ventral stream provides allocentric information to augment this transformation by stabilizing ‘noisy’ egocentric signals [17,20]. But again, direct causal evidence to link these observations to the spatial deficits observed in patient populations is lacking. One potential approach is to transiently perturb ongoing neural processes with transcranial magnetic stimulation (TMS). For instance, TMS over known parietal ‘reach regions’ has been shown to reduce egocentric precision and accuracy in memory-guided reaches [23–26]. One previous TMS study showed that stimulation of both the ventral and dorsal streams can influence movement time in an allocentric reach task [27]. However, to our knowledge, no previous TMS study has provided the dissociation necessary to directly establish causal evidence for the dorsal / ventral segregation of egocentric / allocentric coding of reach targets.

We tested this question by applying online TMS to MRI-localized occipital hubs identified in our previous study [8] during the memory interval before reaches, using a previously reported psychophysical paradigm [5]. Based on the observation that allocentric information is integrated into dorsal stream mechanisms [20,17,5], we predicted that superior occipital stimulation would affect both egocentric- and allocentric-based reaching, inferior occipital stimulation would selectively impair only allocentric performance, and that these effects would likely interact with asymmetries observed in the control data [5]. In addition, we performed a network behavior analysis to determine TMS site influences, based on the modularity of subnetworks derived from individual resting state MRI data and reach errors. Consistent with our hypothesis, superior occipital stimulation perturbed behaviour in both tasks, inferior stimulation only perturbed behaviour in the allocentric task, and further dissociations between dorsal and ventral spatial coding merged in detailed behavioral asymmetries and cross-participant modularity–behaviour correlations.

## Results

### Overview of Experimental Paradigm

Based on an *a priori* power analysis, 16 participants performed each stage of the experiment (see Methods for details). Each series of experiments (Fig 1) began with an MRI session designed to localize TMS targets and establish functional network modularity. Fig 1A shows the two stimulation sites in the left inferior (ExtrInf-2) and superior (ExtrSup-2) extrastriate occipital cortex (i.e., contralateral to the right hand used in the reach task) [28], superimposed on group-average population receptive field maps. In our previous functional network analysis, these were identified as important hubs in an inferior occipital-temporal and superior occipital-parietal module, respectively [8]. Henceforth we refer to these as the inferior occipital (InfOcc-TMS) and superior occipital (SupOcc-TMS) stimulation sites.

**Fig 1.**
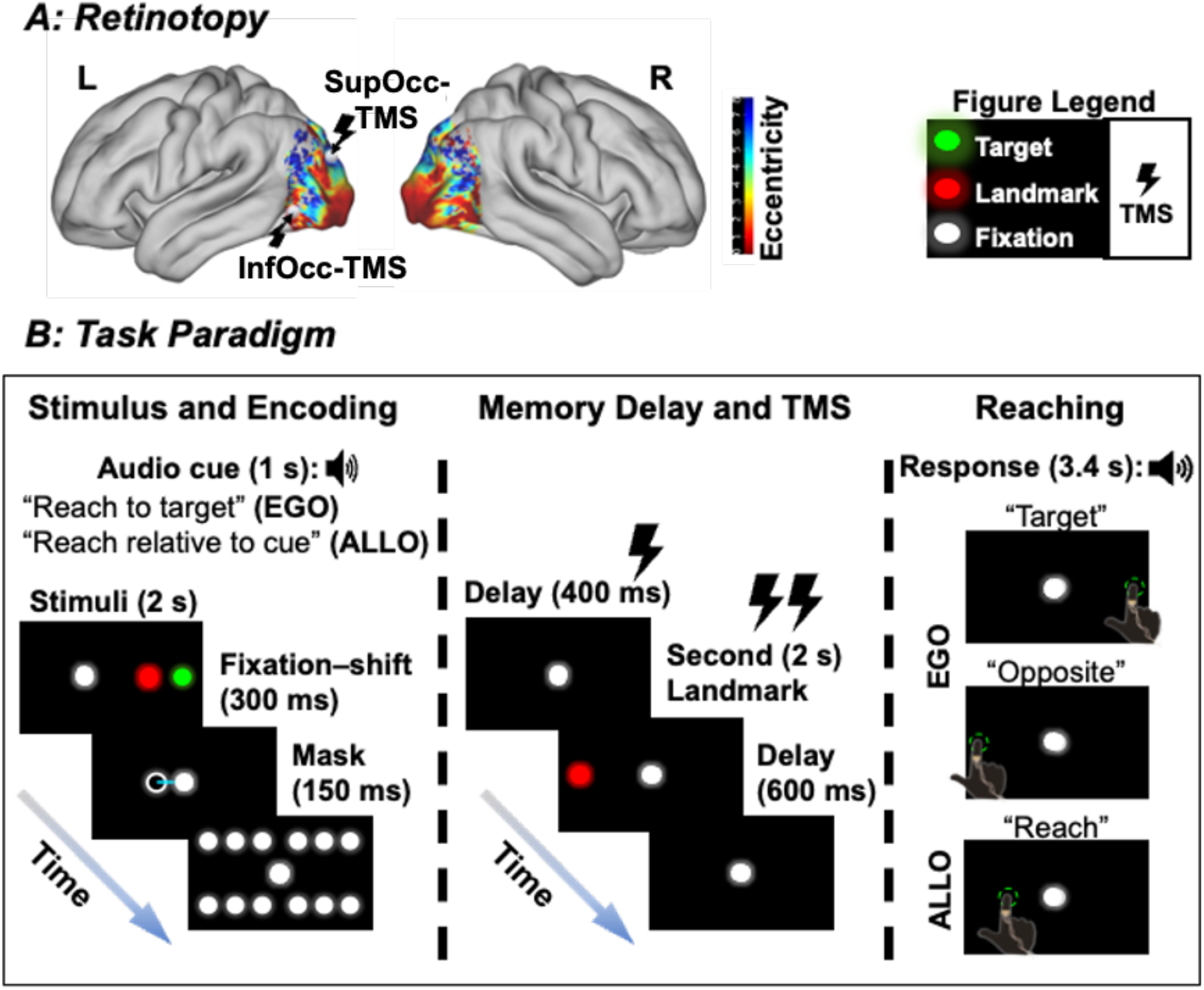
Stimulation site and experimental paradigm. **A –** TMS sites (SupOcc-TMS and InfOcc-TMS) superimposed on group-average Population Receptive Field Model maps of target eccentricity (N = 16) are shown on the left and right lateral cortex. Colors indicate eccentricity according to the color scale, showing central (red) to peripheral (blue) eccentricity. **B –** Experimental paradigm summarized across three task epochs. **Stimulus and encoding:** Participants fixated a white dot while a landmark (red dot) and target (green dot) briefly appeared. Fixation then shifted to the center to dissociate egocentric (EGO) and allocentric (ALLO) reference frames, followed by a mask. **Memory delay and TMS:** TMS (three pulses at 1 Hz) was delivered during the first delay and when the landmark reappeared at a different position. **Reaching:** An auditory cue instructed participants to reach and touch with their right index finger the remembered target location according to either the EGO instructions (“Target”/”Opposite”) or ALLO instructions (“Reach”).

During behavioral / TMS experiments (Fig 1B), participants were instructed at the start of each trial to either ‘reach to target’ (EGO task) or ‘reach relative to cue’ (ALLO task). Participants then fixated a white dot while a landmark (red) and target (green) briefly appeared within a horizontal stimulus array (encoding phase). Fixation subsequently shifted to the centre to dissociate egocentric and allocentric reference frames, followed by a visual mask. This was followed by the memory delay, during which the landmark re-appeared at a different horizontal position in the same or opposite visual field.

Finally, an auditory instruction cued a reach with the right index finger to touch the remembered target location. In the ALLO task, the instruction was simply: ‘reach’ This could result in reaches toward the same visual field (ALLO-SAME task) or opposite visual field (ALLO-OPP task) as the original target, depending on where the landmark reappeared. To balance this, we used two instructions in the EGO task: toward the “Target” (EGO-SAME task) or “Opposite” (EGO-OPP task). Note that visual stimuli were the same in both tasks, only the instructions were different. Stimulation trials were the same, except three TMS pulses were applied at 1 Hz during the memory delay, spanning landmark re-appearance.

### Reach Endpoint Errors

Fig 2 illustrates how we quantified reach errors. The top two panels show target locations (black dots labeled -12° to +12°). All panels show the corresponding participant average reach end-point positions per target location (coloured dots), and 95% confidence ellipses fit to the reach data, averaged across participants. The centres of these ellipses correspond to the mean constant error for each target and their area corresponds to the mean variable error.

**Fig 2.**
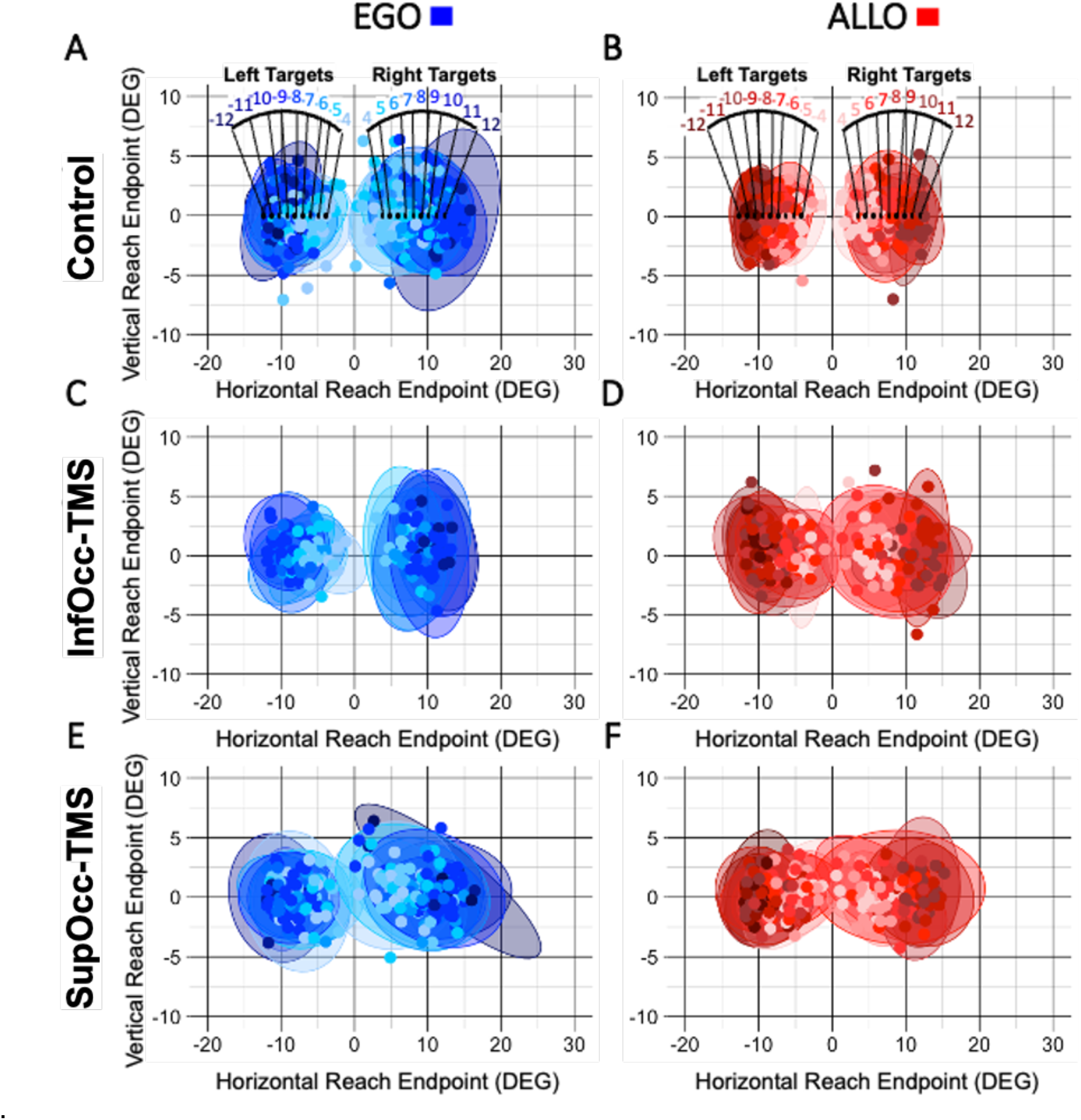
Average Reach Endpoint Ellipses. Average 95 % confidence ellipses (2D reach endpoint variability) at each target location are shown for EGO (blue) and ALLO (red) tasks and across stimulation conditions: **A,B –** Control data, including target positions in visual angle degrees as black dots numbered 4 - 12 relative to central fixation right and left (negative values relative to fixation) visual fields. **C,D –** InfOcc-TMS, and **E,F –** SupOcc-TMS. Small, filled circles are individual participant average reach endpoints. Light-to-dark colors represent data for targets running central to peripheral. SAME / OPP side trials are pooled in this dataset.

Fig 2 panels A and B illustrate the control condition for EGO (blue) and ALLO (red) tasks, respectively. Reach distributions clustered around the correct target locations, i.e., ellipses shift from left to right along with the targets. Further, the EGO ellipses are visibly larger (mean 18.64 ± 5.25 deg²) than the ALLO ellipses (15.92 ± 5.05 deg²). This replicates our previous finding that an allocentric instruction leads to more precise overall performance in memory-guided reaching tasks [5].

Reach performance looks generally similar during TMS trials (Fig 2 C-F) but with subtle differences that are already evident in the raw data. For example, the allocentric advantage observed in control trials (Fig 2 B) appears to be lost during both InfOcc-TMS (Fig 2 D) and SupOcc-TMS trials (Fig 2 F). These and other observations are documented and quantified more extensively in the following sections and figures.

### Influence of TMS in the Spatial Domain: Variable and Constant Error

Our hypothesis predicts that SupOcc-TMS will impair both egocentric and allocentric pointing (because they share a common sensorimotor mechanism), whereas InfOcc-TMS will affect only allocentric reaching. Fig 3 A and B compare spatial performance in our different experimental conditions by providing the distributions of variable error (mean ellipse areas) and constant error (mean absolute error) across participants, including both LEFT/RIGHT field and SAME/OPP reaching for ALLO and EGO conditions.

**Fig 3.**
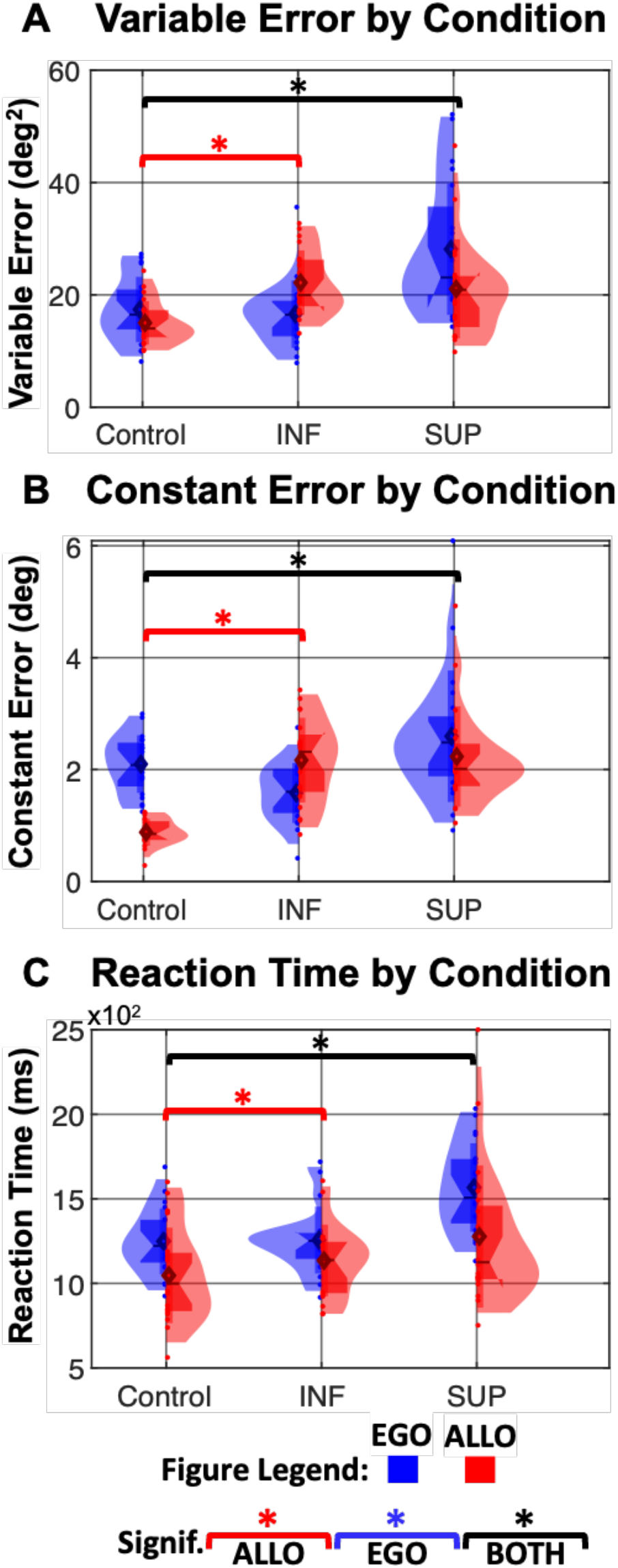
Influence of TMS in the Spatial and Temporal Domains. Violin plots showing the distribution of (top) **A –** Variable Error (deg²), (middle) **B –** Constant Error (deg), (bottom) **C –** Reaction Time (ms), for EGO (blue) and ALLO (red) conditions across Control, InfOcc-TMS (INF), and SupOcc-TMS (SUP) conditions. Horizontal lines indicate significant differences (p < 0.05), color coded by condition. LEFT / RIGHT field and SAME / OPPOSITE reaching trials are pooled in these datasets but are separated in Supplementary Figure 1.

First, we examined the influence of TMS on variable errors relative to control performance (Fig 3 A). A mixed-effects model revealed a significant interaction between stimulation site and task condition (interaction: *t* = 6.21, *p* = 5.4 × 10⁻¹⁰), indicating that the effect of TMS differed between EGO and ALLO trials. Follow-up one-tailed pairwise t-tests for increased error of the interaction effect (adjusted for multiple comparisons) showed that SupOcc-TMS increased reach variability in both EGO (Δ 9.52 deg²; *t* = 10.62, *p* < 1 × 10⁻¹⁶) and ALLO trials (Δ 4.89 deg²; *t* = 4.66, *p* = 1.79 × 10⁻⁶) relative to control. In contrast, InfOcc-TMS stimulation produced a selective effect, with no reliable change in EGO trials (*t* = 1.21, *p* = 0.12), but a significant increase in ALLO variability (Δ 3.57 deg²; *t* = 3.08, *p* = 1.06 × 10⁻³).

Next, we quantified constant error as the mean absolute reach error between the goal and the actual reach (Fig 3 B). A mixed-effects model again revealed a significant interaction between stimulation site and condition (*t* = 7.48, *p* = 8.9 × 10⁻¹³), indicating differential TMS effects across tasks. In the control condition, mean error was higher for EGO trials (2.02°) than ALLO trials (1.07°), replicating Musa et al. [5]. SupOcc-TMS increased constant error in both EGO (Δ 0.87°; *t* = 5.63, *p* < 1 × 10⁻¹⁶) and ALLO trials (Δ 1.19°; *t* = 11.92, *p* < 1 × 10⁻¹⁶). In contrast, behaviour during InfOcc-TMS stimulation showed no significant increase in EGO trials (Δ −0.77°; *t* = −3.95, *p* = 0.99) while increasing error in ALLO trials (Δ 1.27°; *t* = 6.56, *p* < 2 × 10⁻¹⁶).

In summary, the influence of TMS in the spatial domain depended on both stimulation site and task: TMS over superior occipital cortex influenced spatial reach errors in both tasks, but TMS over inferior occipital cortex only affected the ALLO task.

### Influence of TMS in the Temporal Domain: Reaction Time

Assuming that our TMS pulses interfered with motor planning, our hypothesis remains similar for reaction time: deficits in both tasks during SupOcc-TMS, but only the ALLO task during InfOcc-TMS. Temporal results are shown on the bottom panel of Fig 3 C. A mixed-effects model revealed a significant interaction between stimulation site and task condition (*t* = 5.84, *p* = 4.7 × 10⁻⁹), indicating that the effect of TMS differed between EGO and ALLO trials. In the control condition, reaction times were longer for EGO trials (mean: 1211 ms) than for ALLO trials (1017 ms), replicating findings from Musa et al. [5]. Follow-up one-tailed pairwise contrasts testing an increase in reaction time showed that SupOcc-TMS significantly prolonged reaction times in both EGO (Δ 359 ms; *t* = 6.29, *p* < 1 × 10⁻¹⁶) and ALLO trials (Δ 373 ms; *t* = 6.91, *p* < 1 × 10⁻¹⁶). In contrast, InfOcc-TMS increased reaction times in ALLO trials (Δ 76.3 ms; *t* = 3.08, *p* = 0.0011), but did not reliably affect EGO trials (*t* = 1.02, *p* = 0.16).

In summary, the influence of TMS in the temporal domain depended on both stimulation site and task. TMS over superior occipital cortex influenced reaction time in both tasks, but TMS over inferior occipital cortex only affected the ALLO task. Taken together with our spatial results (Fig 3 A,B), these results support our hypothesis that the superior occipital hub is part of the default sensorimotor transformation for reach, whereas the inferior occipital hub is only engaged for allocentric processing.

### Spatial Asymmetries: Visual Field and SAME vs. OPPOSITE reaching

The reach system tends to show contralateral preference for both hand and visual field [30,31]. Based on this, we expected that 1) left TMS would have the most influence for reaches in the right visual field, 2) this effect would be greater in the ALLO task during InfOcc-TMS, and 3) less task-dependent during SupOcc-Stim. Consistent with this, we found an interaction between task, stimulation site, and visual field asymmetries (Figure 4A/B; t = 3.99, p = 6.8 × 10^−5^). Inferior stimulation increased RIGHT > LEFT asymmetry for both variable error (Figure 4A, Δ = 2.93°; t = 3.72, p = 2 × 10^−4^) and constant error (Figure 4B, Δ = 2.27°; t = 6.56, p = 5 × 10^−11^) in the ALLO task (red), with no (or decreased) asymmetry in the EGO task. Superior stimulation increased variable error in the EGO task (Figure 4A, Δ = 1.98°; t = 4.56, p = 5 × 10^−6^) and constant error in both tasks (Figure 4B, ALLO task: Δ = 1.24°; t = 2.56, p = 0.01; EGO task: Δ = 2.06°; t = 4.01, p = 6 × 10^−5^).

**Fig 4.**
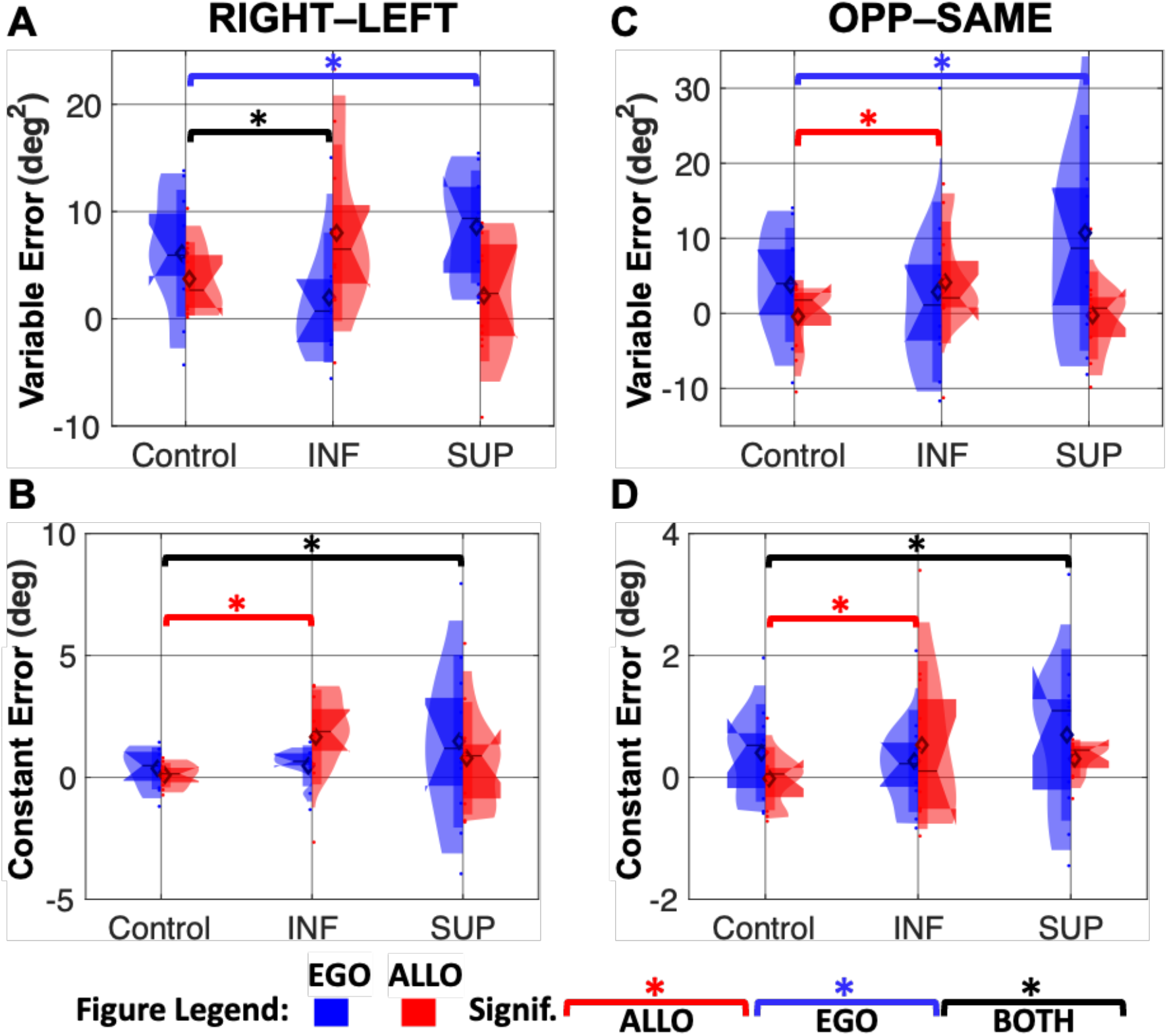
Lateralization of Response. The average **Precision** and **Accuracy** compared for **A,B –** OPPOSITE-SAME target-movement mappings and **C,D –** RIGHT-LEFT visual field lateralization for EGO (blue) and ALLO (red) conditions across Control, TMS InfOcc-TMS, and TMS SupOcc-TMS conditions. Horizontal lines indicate significant differences (p < 0.05), color coded by condition.

In our previous study, we found that egocentric reach performance was less accurate and precise when the movement goal appeared in the opposite visual field to the target, whereas the presence of a landmark in the ALLO task largely nullified this effect [5]. This was confirmed in our control data (Fig 4 C/D). If TMS affects this OPPOSITE > SAME error pattern, our hypothesis would predict this influence to be task dependent: inferior occipital stimulation for the ALLO task, superior occipital stimulation for both tasks. Consistent with this, we found a significant interaction between TMS stimulation site and task in these data (Figure 4 C/D; t = 5.84, p = 4.7 × 10⁻⁹). InfOcc-TMS re-introduced OPPOSITE-SAME asymmetries in the ALLO task for variable (Fig 4C; Δ = 3.23°; t = 2.83, p = 0.0046) and constant (Fig 4D; Δ = 1.19°; t = 2.07, p = 0.038) errors, whereas superior stimulation exacerbated the pre-existing egocentric asymmetries for variable error in the EGO task (Fig 4C; Δ = 6.93°; t = 2.83, p = 0.0046) and constant error (Fig 4D; Δ = 1.45°; t = 2.03, p = 0.042).

In summary, the influence of TMS on visual field and same-opposite goal asymmetries was site-dependent and generally consistent with our hypothesis: inferior occipital stimulation affected both variable and absolute errors in the ALLO task, whereas superior stimulation affected variable errors in the EGO task and constant errors in both tasks. Moreover, TMS tended to exacerbate errors associated with dissociating the movement goal from the target, either negating the advantage of a landmark (during inferior stimulation) or magnifying control errors (during superior stimulation). Finally, similar TMS asymmetries were also observed in reaction time (Supplementary Fig 2).

### Network Analysis: Modularity Predicts Behavioural Changes Induced by TMS

It is thought that TMS has an influence well beyond local networks [32,32]. To test the relationship between TMS site, regional network properties, and behavior, we performed a functional network analysis based on our resting state MRI data [29]. Specifically, we segregated the imaging data of each participant into 8 bilateral network subdivisions (Fig 5 A), corresponding to the Control, Default, Dorsal Attention, Limbic, Visual, Somato-Motor, Salience Ventral Attention and Tempero-Parietal Networks [28]. We then used graph theory analysis to compute the modularity of each subdivision (the degree to which network nodes tend to cluster into correlated communities). We found that, across participants, the modularity score for most of the brain subdivisions listed above were significantly correlated to both constant and variable behavioral errors measured in our control data (Supplementary Fig 3).

**Fig 5.**
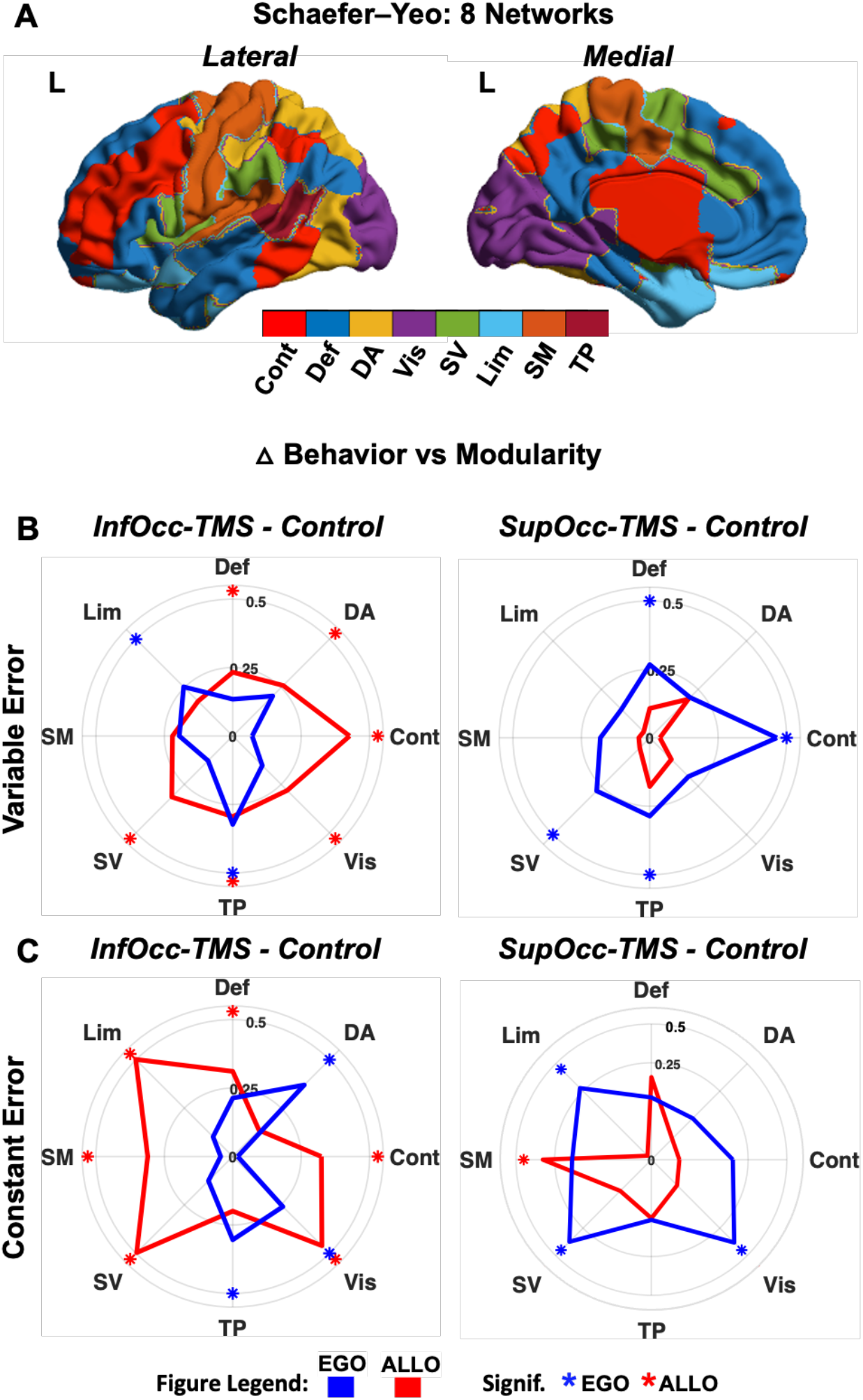
Association of Network Modularity with Behaviour. **A –** Schaefer-Yeo 8 network parcellation (collapsed from 17 networks to 8 unique networks) is color-coded by network and overlaid on the left medial and lateral cortex. The 8 parcellations are: Control (Cont), Default (Def), Dorsal Attention (DA), Limbic (Lim), Visual (Vis), Somato-motor (SM), Salience Ventral Attention (SV) and Tempero-parietal (TP) Networks. The *four lower panels* **(B, C)** show the relationship between modularity and TMS-induced **B –** variable and **C –** absolute errors in the 8 networks described above, plotted radially. The *left column* plots InfOcc-TMS data and the *right column* plots SupOcc-TMS data, during the EGO (blue) and ALLO (red) tasks. Each datapoint represents the relationship (slope) between modularity and TMS-induced errors across 16 participants (quantified as the absolute value of the TMS induced error vs modularity linear association), computed using standardized regression coefficients. Asterisks indicate statistical significance.

We then tested if modularity predicts TMS-induced errors (i.e., TMS - control error), the cortical distribution of this influence, and its dependence on TMS site. Fig 5 B/C show the results of this analysis for variable (B) and constant (C) error parameters. Each radial datapoint represents the relationship (regression slope) of TMS-induced errors as a function of modularity across 16 participants, computed for each of the 8 brain subdivisions. Both error measures show that 1) modularity predicts TMS-induced error across a broad range of cortical subnetworks (with strongest associations in the Visual, Tempero-Parietal, Control and Salience Ventral Attention networks), and 2) although significant modularity-behavior influences were present during stimulation of both sites and tasks, InfOcc-TMS had more influence in the ALLO task (red) and SupOcc-TMS had more influence in the EGO task (blue). Taken together with our other results, this suggests that (although the dorsal stream participates in both tasks), its intrinsic network organization is primarily egocentric.

## Discussion

The present study used transcranial magnetic stimulation (TMS) to investigate the causal role of extrastriate visual cortex in egocentric (EGO) and allocentric (ALLO) memory-guided reaching. We hypothesized that superior occipital stimulation would affect both egocentric- and allocentric-based reaching, whereas inferior occipital stimulation would selectively impair allocentric performance. Consistent with our hypotheses, stimulation of the superior occipital cortex disrupted performance (accuracy, precision, and reaction time) in both EGO and ALLO conditions, whereas stimulation of the inferior occipital cortex produced selective impairments that were strongest in the ALLO condition. TMS also had more specific effects, i.e., tending to affect movements to contralateral space and target-goal dissociations (in the OPPOSITE condition), in the predicted site and task-dependent manner. Finally, a double dissociation between superior / EGO vs. inferior / ALLO emerged in the widespread relationship between occipital stimulation, cortical modularity, and behavioral errors. Together, these findings consistently show a causal dissociation between the contributions of inferior and superior occipital cortex to spatial behavior.

### Behaviour in the Control Task Replicates Previous Observations

Even in the absence of TMS, the instruction to use EGO vs. ALLO cues influenced reaching behaviour. ALLO reaching was more precise and accurate than EGO reaching, as reflected by smaller endpoint variability and lower absolute error. Reaction times were also faster in the ALLO condition, indicating more efficient movement planning when landmark information was available. These findings replicate the results of our previous behavioral study [5], and support the idea that allocentric landmarks help stabilize spatial behavior in the presence of spatial uncertainty and ‘noisy’ internal egocentric signals during memory delays [13,17,34].

### TMS Causally Dissociates Visual Streams and Spatial Behavior

As noted in the introduction, neuropsychological, neuroimaging, and neurophysiological data suggest that the ventral visual stream is associated with allocentric spatial processes but ultimately must influence behavior through a primarily egocentric dorsal stream [4,6–8,20,21]. But, causal evidence for this dissociation in healthy humans was lacking until now. This led us to hypothesize that inferior occipital TMS would selectively influence landmark-guided reaching, whereas superior occipital TMS would influence both allocentric and egocentric reaching. Overall, our TMS results (selective disruption of reach precision, accuracy, and reaction time) support this prediction. By design, the timing of our TMS pulses (during the memory interval preceding action), suggests that these errors arise from disruptions of spatial memory and planning [23], as opposed to feedback control during execution [25,26].

These results are consistent with the numerous studies which suggest transformation from gaze-centred visual signals to somatotopic motor signals through the dorsal occipital-parietal-frontal pathway [10,18,19,35,36] in contrast to configurational (feature relative to feature, object relative to object) processing in the ventral stream [10,11,37]. However, this distinction is not absolute: allocentric relationships (such as target-relative-to-landmark) must ultimately be represented in some sensory frame (such as the eye) and be converted into muscle coordinates to influence behavior. Behavioral studies suggest that this transformation occurs at the first opportunity, possibly to benefit readiness for action [20]. This is likely why allocentric cues influence the dorsal stream [20], even extending to single-cell codes in frontal cortex [39].

### Occipital TMS Influences Spatial Asymmetries in a Site-Specific Manner

Focal TMS in the visuomotor reach network is known to have more influence on the contralateral visual field and limb [23,24,39–41] and has been shown to disrupt the stability of location when eye movements dissociate the target from the movement goal [42,43], especially when the goal is updated into the stimulated hemifield [44]. A similar spatial updating process occurs in ‘anti-reaching’ tasks, like the EGO-OPPOSITE task used here [1,45,46]. As noted above, these processes are known to benefit from the stabilizing influence of visual landmarks [3,5].

Our TMS results show that the stimulation site interacts with these spatial asymmetries in a site-dependent manner consistent with our main hypothesis: first, we observed that inferior occipital stimulation had more contralateral influence in the ALLO task, whereas superior stimulation was more task-independent. Second, we observed that inferior stimulation tends to ‘undo’ the benefit of landmarks in the OPPOSITE task, whereas superior stimulation tends to exacerbate the normal errors.

### Implications for Cortical Network Organization

Our findings support a network perspective of visual function in two ways. First, our stimulation sites were derived from a specific inferior occipital hub (for the ALLO task) and superior occipital hub (for both the EGO and ALLO tasks) derived in our previous functional network study [8]. The current results thus provide causal evidence to support the previous study. Second, the new network analysis here establishes a relationship between occipital TMS site, modularity throughout the cortex (based on resting state data), and behavioral errors.

The latter analysis provided several useful observations. First, our control results (Supplementary Fig 3) support the notion that modularity provides a useful biomarker for linking intrinsic network properties to task and behavior [8,29,47]. Second, we found that modularity throughout the brain predicted TMS-induced behavioral errors, perhaps through increased susceptibility to TMS, or by reducing intermodular communication [48–50]. Third, these effects were site dependent in a manner generally supportive of our dorsal-ventral hypothesis. However, the dissociation was stronger than expected (there was a double dissociation — stronger inferior site predictions in the ALLO task, stronger superior site prediction in the EGO task), perhaps because this network-based analysis better reflected the internal organization of the dorsal-ventral streams than overt behavior alone. Finally, this analysis gives general causal credence to the motion that functional network parameters provide real interpretive power for understanding brain-behavior relationships [22,41,48,49,51,52].

### Potential Clinical Implications

The selective impairment of allocentric processing following inferior occipital stimulation has potential clinical relevance. It is well established that dorsal stream damage produces egocentric deficits, whereas ventral stream damage affects landmark-based spatial abilities [6,7, 53–55]. Our results, which show both regional specializations with widespread network influence, provide causal support for the notion that rehabilitation strategies could leverage alternative networks when default spatial mechanisms are compromised [7]. For example, training on the use of allocentric cues may augment or even bypass egocentric mechanisms that have become ‘noisy’ due to damage, disease or normal ageing [8]. Further, the interaction between hub location and network organization may inform individualized clinical neuromodulation.

## Conclusions

This study demonstrates that occipital visual regions make causal and dissociable contributions to memory-guided reaching. Inferior occipital cortex preferentially supports allocentric representations, whereas superior occipital cortex likely integrates allocentric information but fundamentally supports the default egocentric transformations for action. Critically, the behavioural impact of focal disruption is shaped by large-scale network organization. These findings provide a framework linking global network topology, local cortical specializations, and spatial behaviour.

## Materials and Methods

### Participants

Sixteen individuals (8 males, 8 females; ages 22–34 years) were recruited based on an a priori power analysis (See Sample Size Analysis below.) and provided informed consent to participate in this study. All participants were right-handed, had normal or corrected-to-normal vision, and reported no history of neurological disorders, including epilepsy. Participants met standard safety criteria for transcranial magnetic stimulation [56]. No participant data were excluded from analysis. All received monetary compensation for their time. The experimental procedures were approved by the York Human Participants Review Subcommittee (Certificate #: e2019-248) and were conducted in accordance with the tenets of the Declaration of Helsinki.

### Experimental Design

The experiment was conducted across three sessions: one MRI session followed by two TMS sessions scheduled 3 days to 1 week apart. Each session included a control block (no stimulation; 72 trials) with EGO and ALLO trials pseudo randomly interleaved as in Musa et al. [5], followed by a TMS block (72 trials) using the same task structure. The stimulated brain region was counterbalanced across days. Stimulation sites (Fig 6A) were selected based on functional hubs identified in Musa et al. [8]: ExstrSup-2, a superior occipital hub shared by both EGO and ALLO reaching, and ExstrInf-2, an inferior occipital hub preferentially associated with ALLO reaching.

**Fig 6.**
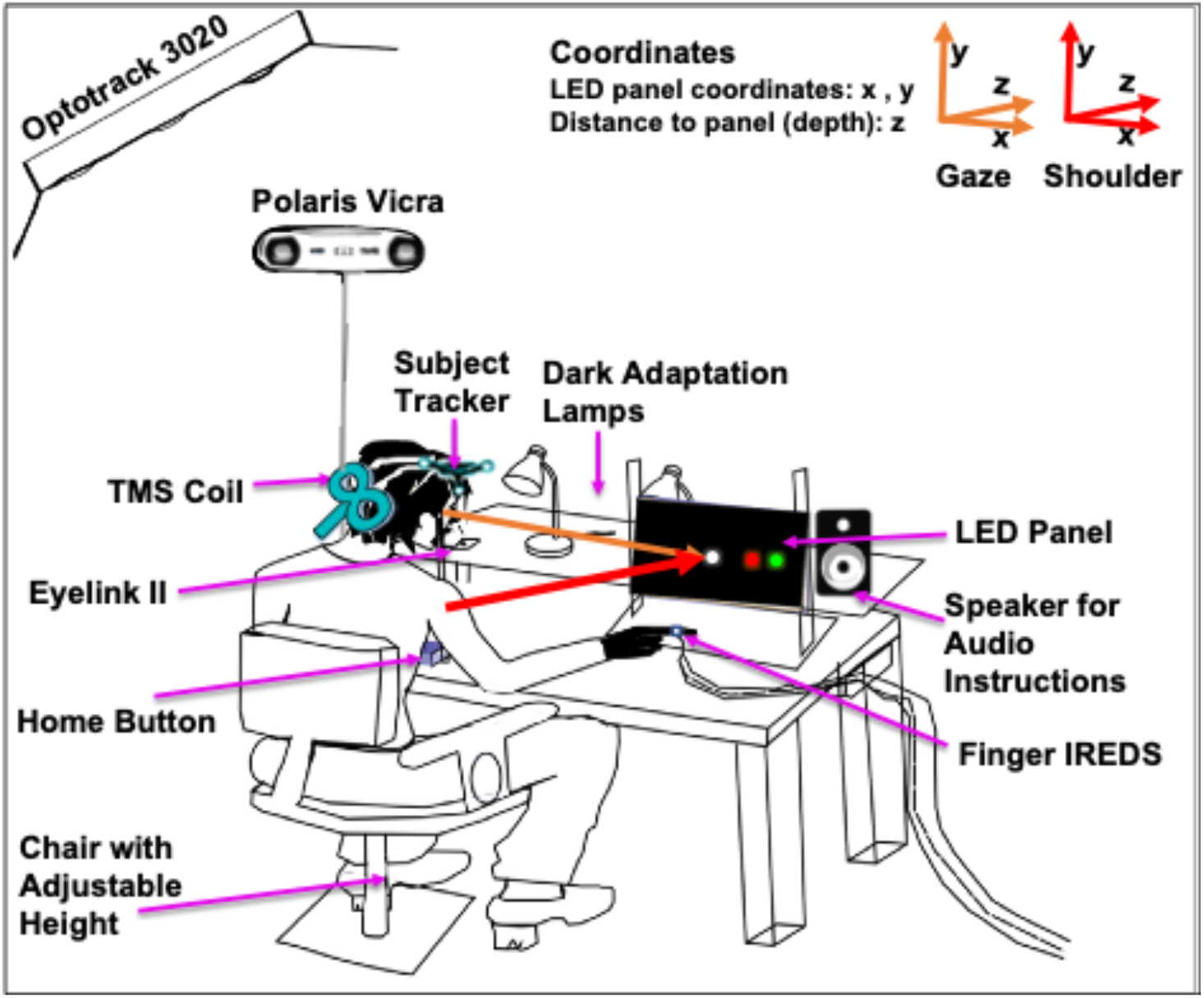
Experimental Setup. Sixteen participants performed a delayed reach-to-touch task under EGO (relative to self) and ALLO (relative to a visual landmark) instructions, with a similar setup to Musa et al., 2024. Participants were seated in an adjustable chair facing an LED panel that presented visual targets. Right index finger position was recorded using infrared markers (finger IREDs) tracked by a motion capture system (Optotrak 3020). Eye position was monitored with an Eyelink II eye tracker. Participants began each trial with their hand on a home button that also controlled the experiment’s pace and executed reaching movements toward targets displayed on the LED panel. Lamps controlled ambient lighting conditions to prevent dark adaptation, and speakers delivered auditory instructions to guide participants during the task. The LED panel was shifted to the right shoulder and coordinate systems were aligned with the center of the LED panel (x–y position on the panel and z depth), with reference frames illustrated for gaze-centered and shoulder-centered coordinates. A TMS coil was positioned over the participant’s head during the stimulation conditions and neuronavigated using a subject tracker attached to the head and a Polaris Vicra camera. TMS (triple-pulse, 1 Hz, 110% resting motor threshold) was applied during the delay + planning period to dorsal / ventral hubs.

### Experimental Setup for TMS and Behavioral Recordings

Participants performed a delayed reach-to-touch task under egocentric (EGO) and allocentric (ALLO) instructions, with sensory and motor demands identical to those of Musa et al. [5]. The full experimental setup is shown in Fig 6. Participants were seated in a darkened room with their heads stabilized using a personalized bite bar. The right hand rested on a button box that served as both the starting position and pacing control. A customized ring with a 3 × 3 array of infrared-emitting diodes (IREDs) was attached to the right index finger, and three-dimensional hand position was recorded using two OptoTrak 3020 tracking systems (Northern Digital, Waterloo, ON, Canada). Eye position was monitored monocularly (right eye) using an EyeLink II infrared eye-tracking system (SR Research, Ottawa, ON, Canada) mounted on the bite-bar stand. Visual stimuli were presented on an LED display panel positioned approximately 50 cm from the eyes and aligned with the participant’s right shoulder, ensuring stimuli were centered within the comfortable mechanical workspace of the right arm. Occasional illumination was provided during breaks to prevent dark adaptation.

### Task Paradigm

The behavioral paradigm is illustrated in Figure 1B and described in the accompanying text of the Results section. As described in Musa et al. [5], each trial consisted of fixation, brief presentation of a visual target (green LED) and landmark (red LED), followed by a memory delay and a reach-to-touch response. Participants received auditory instructions at trial onset specifying whether to reach based on the remembered target location relative to themselves (EGO) or relative to the landmark (ALLO). In a subset of trials, participants performed mirror-opposite (“anti-reach”) responses, as described previously [45,57].

### Transcranial Magnetic Stimulation and Neuronavigation

TMS was delivered using a Magstim Rapid^2^ stimulator (MagStim, Whitland, UK) and a 70-mm Air Film Coil (air-cooled figure-of-eight coil). Subject-specific functional parcellations derived from resting-state fMRI were used to precisely localize the centroid coordinates of Schaefer atlas parcels corresponding to ExstrSup-2 and ExstrInf-2 for stimulation [28]. For simplicity, the current manuscript refers to these TMS targets as SupOcc-TMS and InfOcc-TMS, reflecting their assignment to the superior occipito-parietal and inferior occipito-temporal modules, respectively, in the network analysis of Musa et al. [8]. These coordinates were individualized for each participant by coregistering the T1-weighted anatomical scan using Brainsight neuronavigation software (Version 2.1.5; Rogue Research, Montreal, Canada) for real-time neuronavigation and accurate coil positioning during the TMS session.

During TMS Blocks, three 1 Hz biphasic single pulses were delivered at suprathreshold (110% of visible resting motor threshold [vRMT]) during the memory delay period (Fig. 1B), spanning the second appearance of the landmark. vRMT was defined as the minimum stimulation intensity required for visible contraction of the first dorsal interosseous in 5 of 10 consecutive trials. The final stimulation intensity was adjusted with consideration of the known differences between visually determined and EMG-based motor thresholds and the associated safety implications of the visual observation method [58]. This protocol was designed to perturb memory and planning signals as well as the integration of the landmark information preceding movement onset, while minimizing sensory side effects and muscle discomfort.

### Behavioral Data Acquisition and Analysis

Finger and eye position data were analyzed offline using custom MATLAB scripts (R2019a). Reach endpoints were converted from OptoTrak (Northern Digital Inc [NDI]) coordinates into screen and visual coordinates using participant-specific calibration mappings. Movement onset and offset were defined using velocity-based thresholds. Endpoint variability was quantified using 95% confidence ellipses, and accuracy was assessed via horizontal overshoot error relative to the expected reach goal. Supplementary Fig 4 illustrates the timing of events for an example participant.

### MRI Data Acquisition

#### Structural Imaging

High-resolution T1-weighted anatomical images were acquired for each participant and used for cortical reconstruction, neuronavigation, and functional coregistration.

#### Retinotopic Mapping

Participants completed retinotopic population receptive field (pRF) mapping to confirm the retinotopic organization of occipital cortex and to visualize stimulation targets relative to visual field representations. These maps were generated using a sweeping bar stimulus from the Human Connectome Project (HCP) visual stimulus set. The bar traversed the visual field in multiple orientations while participants maintained central fixation and performed a fixation task to ensure attentional engagement. pRF mapping followed established methods [59–61]. Voxel-wise responses were modeled using a two-dimensional Gaussian pRF model to estimate preferred visual field location and receptive field size. Retinotopic maps and stimulation sites projected onto the cortical surface are shown in Fig 1A.

#### Resting-State fMRI (Multi-Echo)

Participants also completed multi-echo resting-state fMRI scans. Data were denoised using multi-echo independent component analysis (ME-ICA) to separate BOLD and non-BOLD signal components, improving signal quality for individual-level functional parcellation [62].

#### Individualized Functional Parcellation (GPIP)

Individual T1-weighted anatomical images were processed using the recon-all pipeline in FreeSurfer v6.0.1. Preprocessed and denoised resting-state functional data were coregistered to the anatomical images using bbregister and resampled to the fsaverage5 surface using mri_vol2surf with trilinear interpolation. Surface-based smoothing was applied using a 6-mm full-width at half-maximum kernel with mri_surf2surf. Functional time series were normalized to zero mean and unit variance.

Group Prior Individual Parcellation (GPIP; [63]) was used to generate subject-specific functional parcellations. GPIP was initialized using the 200-parcel, 17-network Schaefer atlas [28], ensuring a common labeling scheme across participants while allowing parcel boundaries to adapt to individual functional connectivity patterns. Parcel homogeneity was computed at each GPIP iteration as the mean pairwise correlation between vertices within each parcel, averaged across parcels. Homogeneity values increased and plateaued prior to the final iteration, indicating stable individualized parcellations. These subject-specific Schaefer parcels were used to localize SupOcc-TMS and InfOcc-TMS targets to guide neuronavigation.

#### Network Modularity Analysis

For each participant, we segregated our imaging data into 8 subdivisions, by simplifying the Schaefer-Yeo 17-network subdivisions (Fig 5 A), which included the Control, Default, Dorsal Attention, Limbic, Visual, Somato-Motor, Salience Ventral Attention and Tempero-Parietal Networks. We first computed inter-regional (parcel) associations of our resting state BOLD time series data and then used graph theoretical analysis to compute the modularity of each network subdivision (grouping together regions of interest into their respective network subdivision). Then we used variations of modularity across participants as a predictor for differences in their behaviour during TMS stimulation, using the spatial behavioural variables described above.

#### Statistical Analysis

Linear mixed-effects models were used to analyze accuracy, reaching variability, and reaction time as a function of task (EGO/ALLO) and stimulation condition (control, SupOcc-TMS, InfOcc-TMS), including all interactions. Spatial asymmetries were analyzed by computing difference scores for visual field (LEFT − RIGHT) and reaching direction (OPPOSITE − SAME), which were subsequently analyzed using separate linear mixed-effects models. Models were fitted using the lme4 package in R [64], with participant included as a random intercept. Estimated marginal means and post hoc pairwise contrasts were computed using the emmeans package, and contrast p-values reported were adjusted for multiple comparisons using the Benjamini–Hochberg false discovery rate procedure.

#### Sample Size Analysis

Sample size was estimated using the simulation-based approach described by Green and MacLeod [65] and implemented in the simr package in R, which performs power analyses for linear mixed-effects models fitted with the lme4 package. Consistent with the approach reported by Musa et al. [5], power was estimated using Monte Carlo simulations based on a pilot dataset comprising three participants and 576 observations across task (EGO/ALLO), stimulation site, and stimulation (control vs. TMS) conditions. A linear mixed-effects model was fitted to the pilot data to estimate the fixed effects and variance components. The estimated effect of interest (β = 0.74) was then used as the basis for the power simulations. Sample size was iteratively increased using simr until the estimated statistical power reached 85%, yielding a required sample size of 15 participants.

## Ethics Approval

Use of brief trains of repetitive transcranial magnetic stimulation (rTMS) to temporarily disrupt cortical function related to spatial memory and eye-hand coordination. Certificate #: e2019-248

## Author Contributions

**LM:** Conceptualization, Data Curation, Formal Analysis, Investigation, Visualization, Writing – Original Draft

**GNL:** Investigation – Data Collection, Writing – Review & Editing

**BS:** Investigation – Data Collection

**AR:** Investigation – Data Collection

**SS:** Software, Methodology

**XY:** Resources, Methodology

**MV:** Methodology, Writing – Review & Editing

**JDC:** Conceptualization, Funding Acquisition, Project Administration, Supervision, Writing – Review & Editing

## Supplementary Materials

**Supplementary Fig 1.**
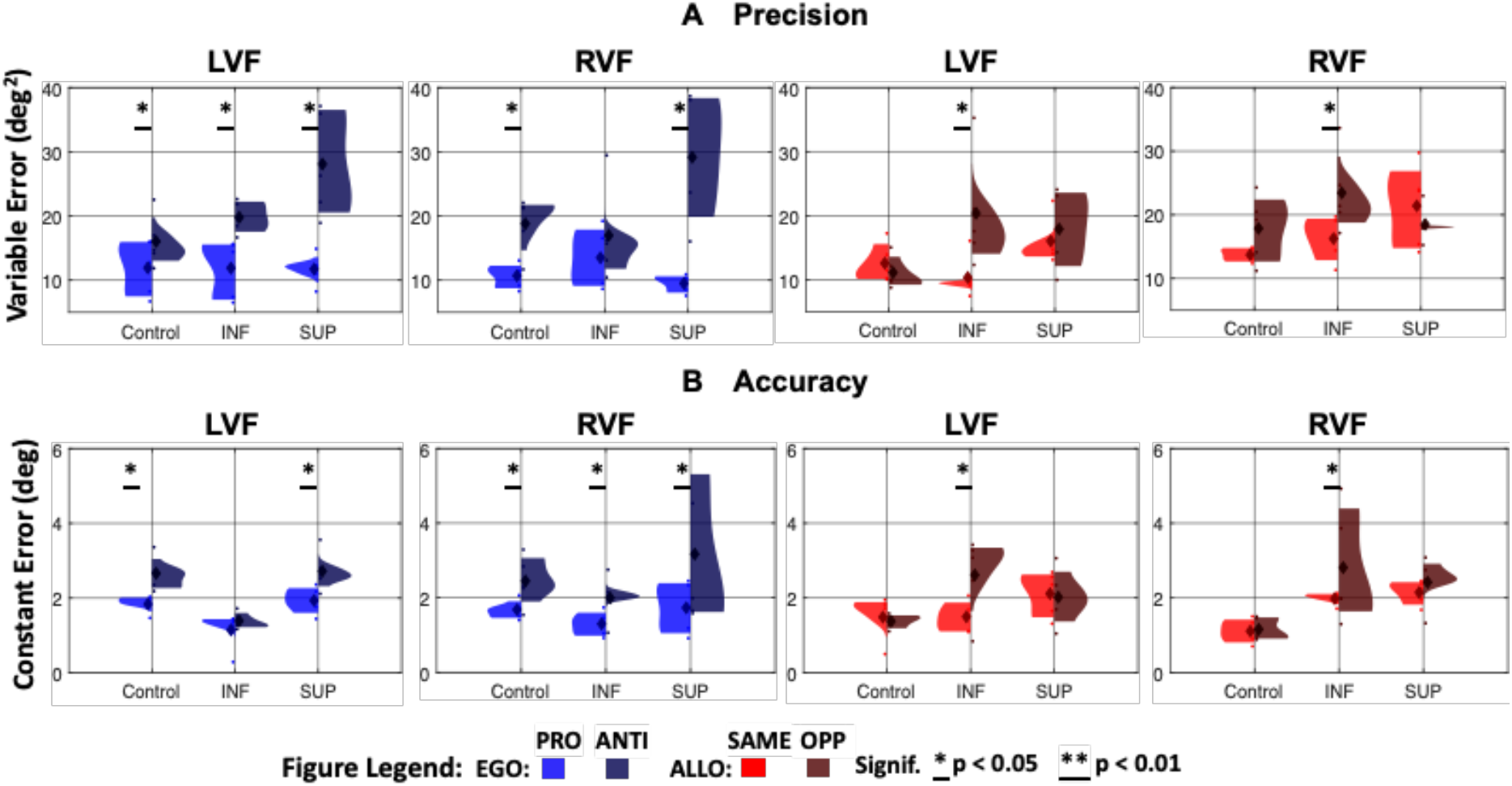
Reach Accuracy and Precision Across Target-Movement Mappings. Violin plots show the distribution of **A – Precision:** Variable Error (deg²) and **B – Accuracy:** Constant Error (deg) for EGO (left panels) SAME/OPP reaching (blue/dark blue) and ALLO (right panels) SAME/OPP second landmark relative to first landmark positions (red/dark red) conditions across Control, InfOcc-TMS, and SupOcc-TMS conditions. Left visual field (LVF) and right visual field (RVF) reaches are shown separately for both spatial updating conditions. Horizontal lines indicate significant differences (p < 0.05).

**Supplementary Fig 2.**
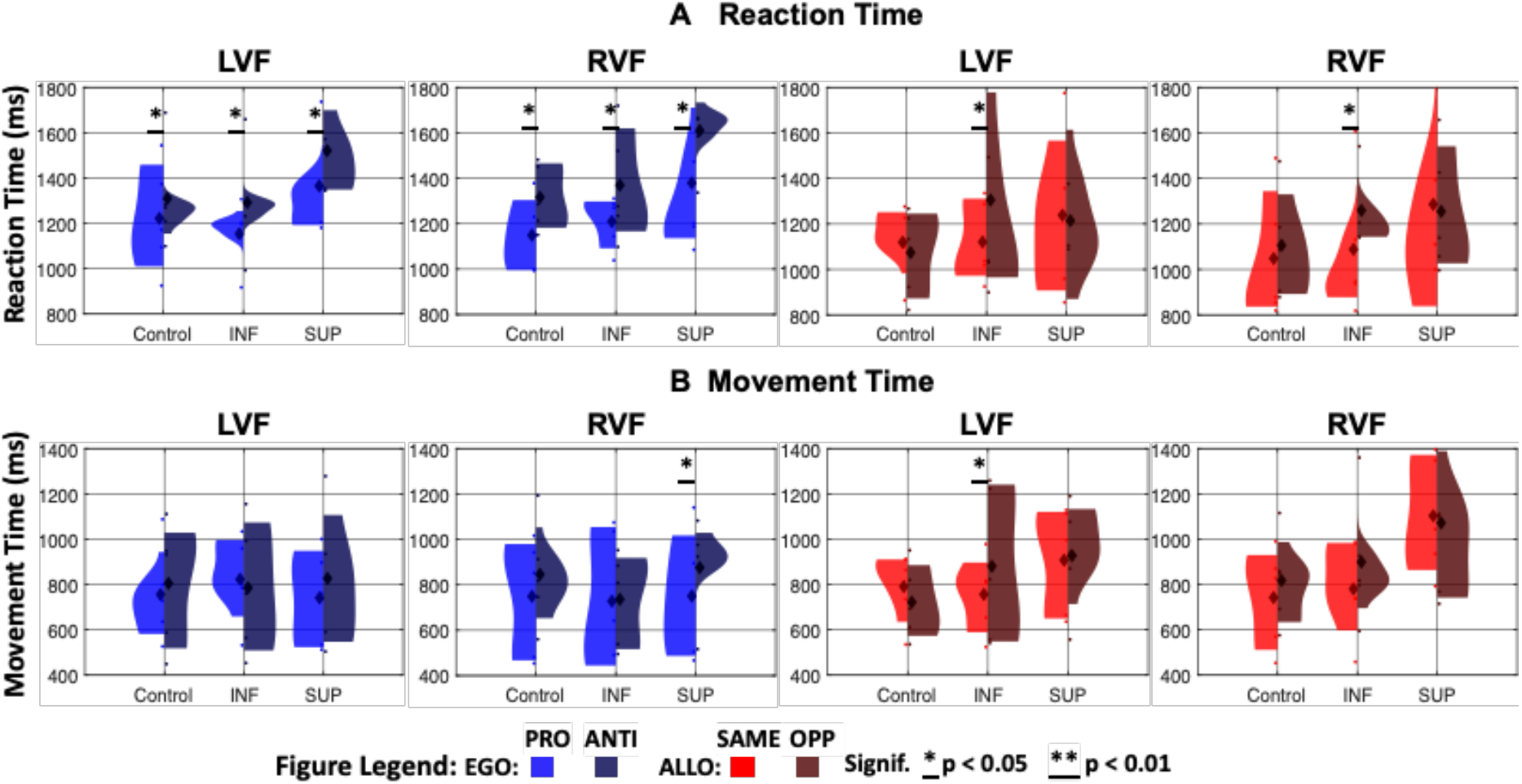
Reaction Time and Movement Time Across Target-Movement Mappings. Violin plots show the distribution of **A – Reaction Time:** Time between Go signal and finger movement onset (ms) **B – Movement Time:** Time between index finger movement onset and stabilization on screen (ms) for EGO (left panels) SAME/OPP reaching (blue/dark blue) and ALLO (right panels) SAME/OPP second landmark relative to first landmark positions (red/dark red) conditions across Control, InfOcc-TMS, and SupOcc-TMS conditions. Left visual field (LVF) and right visual field (RVF) reaches are shown separately for both spatial updating conditions. Horizontal lines indicate significant differences (p < 0.05).

**Supplementary Fig 3.**
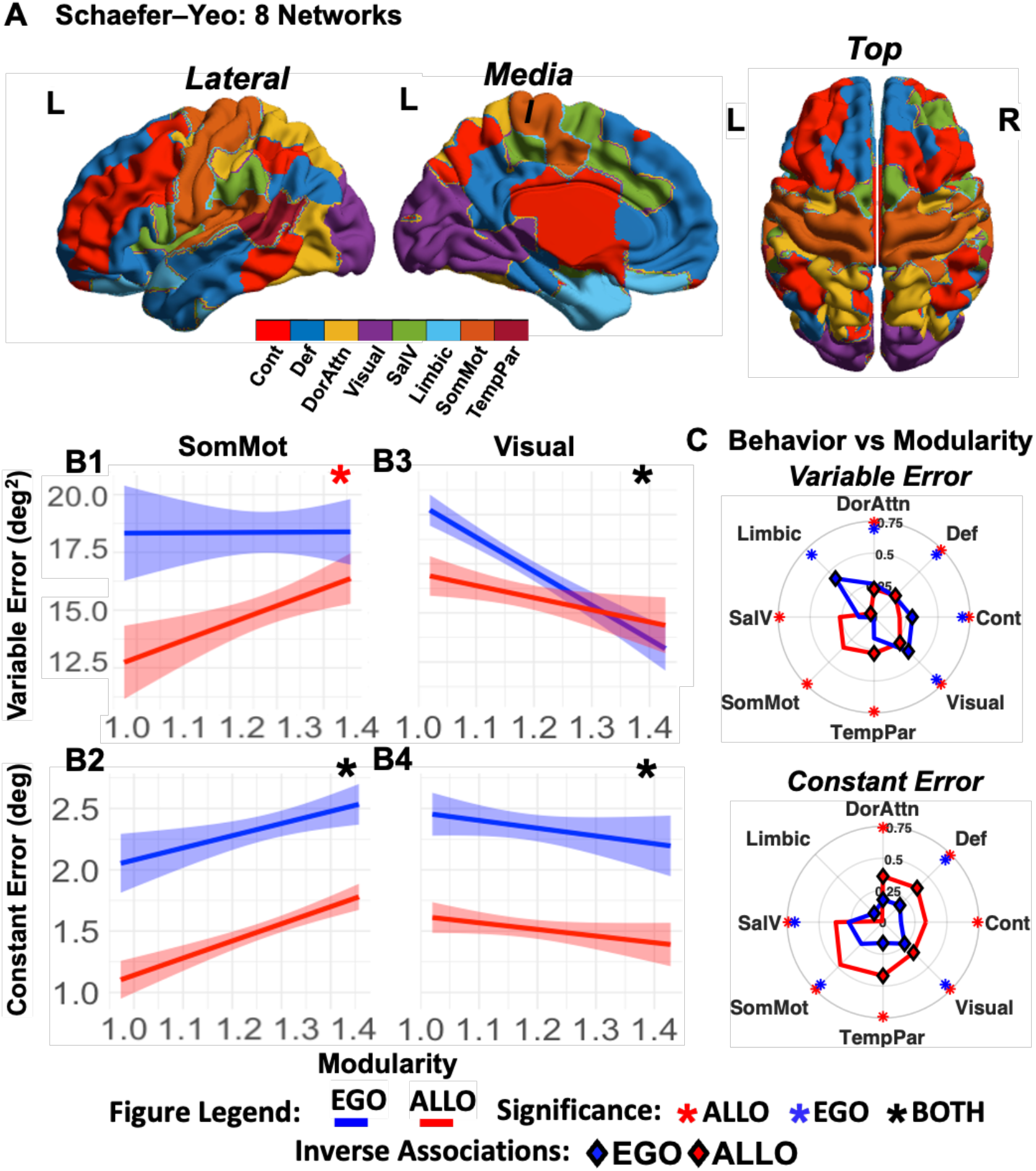
Modularity Correlates to Behavioral Errors in Control Data. **A –** Schaefer-Yeo 8 network parcellation (collapsed from 17 networks to 8 unique networks) is color-coded by network and overlaid on the left lateral cortex. The 8 parcellations are: Control (Cont), Default (Def), Dorsal Attention (DorAttn), Limbic, Visual (Vis), Somato-motor (SomMot), Salience Ventral Attention (SalV) and Tempero-parietal (TempPar) Networks. **B –** The regression plots (line of best fit and 95% confidence interval) on the left show the association between Modularity and Behaviour in the SomMot network (**B1,B2**) and the Visual network (**B3,B4**). Variable Error is shown on the top panels and Constant Error is shown on the bottom panels, for EGO and ALLO conditions in blue and red. and **C –** The two plots in left panel show the relationship between modularity, and variable (top) and constant (bottom) errors for the *Control* condition in the 8 networks described above, plotted radially. Inverse (Negative) associations are indicated by diamond connectors and significant associations are marked in the plot by asterisks, color-coded by condition (blue – EGO, red – ALLO).

**Supplementary Fig 4.**
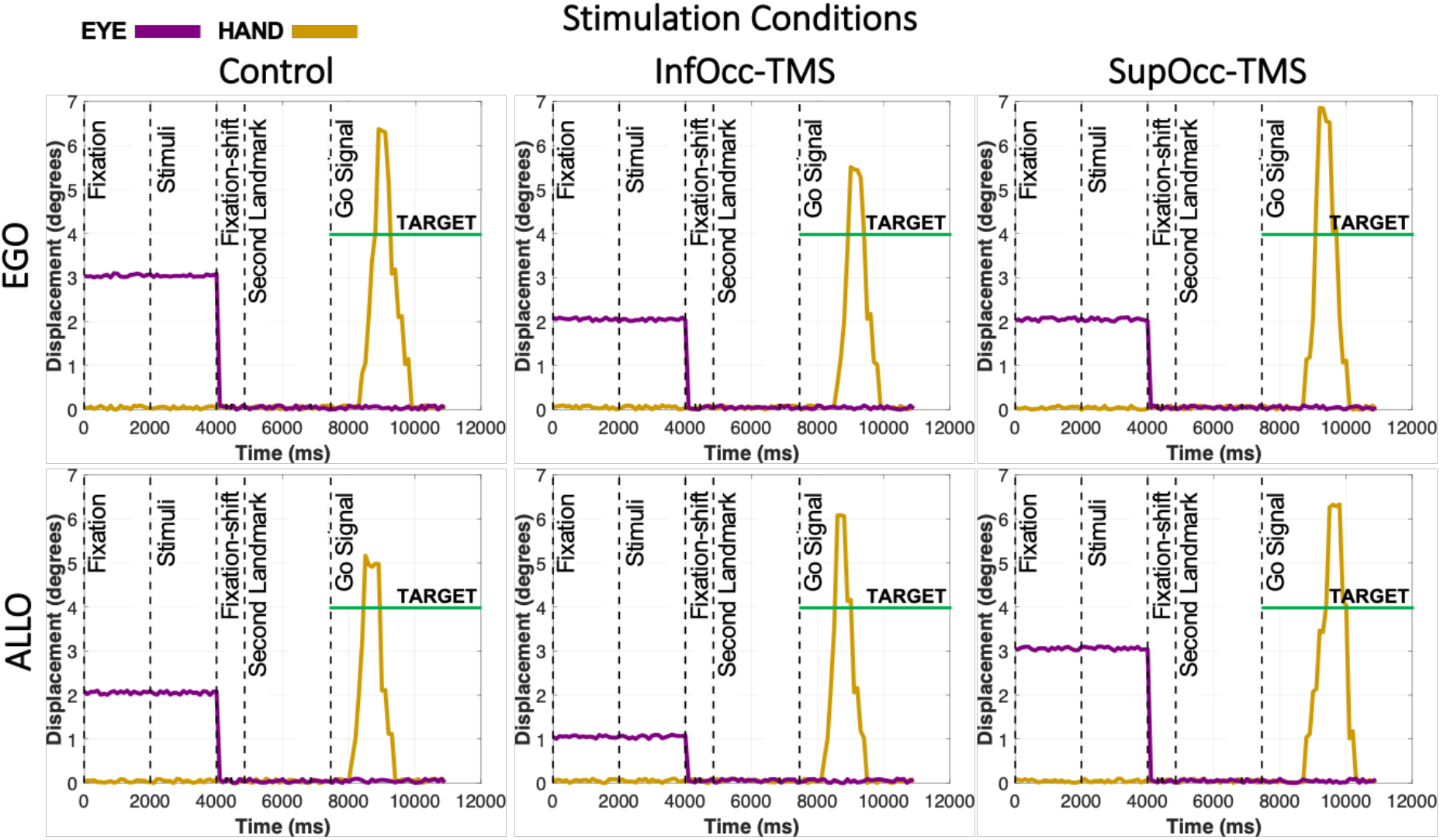
Timing of events. Six example trials from one participant are shown for a target position of 4° under the EGO (top panels) and ALLO (bottom panels) spatial conditions, and for the control and two TMS stimulation conditions. Hand position (purple) and eye position (yellow) are plotted as a function of time. Vertical dashed lines mark the onset of key task events: fixation point appearance (Fixation), target and landmark presentation (Stimuli), and fixation shift to the center of the LED panel (Fixation Shift) during the **Stimulus and Encoding** phase (see Fig. 1 for a description of the task phases and events); landmark reappearance (Second Landmark) during the **Memory Delay and TMS** phase; and the onset of the auditory instruction (Go Signal) during the **Response** phase. The horizontal green dashed line indicates the physical location of the reach target.

